# Tau-first subtype of Alzheimer’s disease consistently identified across in vivo and post mortem studies

**DOI:** 10.1101/2020.12.18.418004

**Authors:** Leon M. Aksman, Neil P. Oxtoby, Marzia A. Scelsi, Peter A. Wijeratne, Alexandra L. Young, Isadora Lopes Alves, Frederik Barkhof, Daniel C. Alexander, Andre Altmann, the ADNI

## Abstract

Alzheimer’s disease (AD) is marked by the spread of misfolded amyloid-β and tau proteins throughout the brain. While it is commonly believed that amyloid-β abnormality drives the cascade of AD pathogenesis, several in vivo and post mortem studies indicate that in some subjects localized tau-based neurofibrillary tangles precede amyloid-β pathology. This suggests that there may be multiple distinct subtypes of protein aggregation pathways within AD, with potentially different demographic, cognitive and comorbidity profiles. We investigated this hypothesis, applying data-driven disease progression subtyping models to post mortem immunohistochemistry and in vivo positron emission tomography (PET) and cerebrospinal fluid (CSF) based measures of protein pathologies in two large observational cohorts. We consistently identified both amyloid-first and tau-first AD subtypes, where tau-first subjects had higher levels of soluble TREM2 compared to amyloid-first subjects. Our work provides insight into AD progression that may be valuable for interventional trials targeting amyloid-β and tau.

## Introduction

Alzheimer’s disease (AD) is a progressive neurodegenerative disease that is characterized at the molecular level by the accumulation of two specific protein-based pathologies within the brain: amyloid plaques, composed of extracellular amyloid-β (Aβ) peptide, and intracellular neurofibrillary tangles (NFTs), composed of abnormally hyperphosphorylated tau protein. These pathologies combine to create a toxic environment that drives neurodegeneration via neuronal and synaptic loss, leading to cognitive impairment^1^. While these protein pathologies have been recognized as the primary signature of AD since Alois Alzheimer first observed them over a hundred years ago^2^, the causal relationship between these pathologies is not fully understood. The prevailing view set forth by the amyloid cascade hypothesis is that the accumulation of Aβ peptides is the main causative event within the pathogenesis of AD, with tau-based NFTs, neurodegeneration and cognitive impairment following as a result^3,4^.

The ‘amyloid-first’ view has strong empirical support in familial AD, where gene mutations (*APP, PSEN1, PSEN2*) associated with abnormal Aβ peptide production have been shown to cause autosomal dominant forms of the disease^5,6^. In contrast, sporadic AD, which accounts for the vast majority of cases, is believed to be caused by a complex combination of genetic and environmental factors^7^. Despite this important difference, the amyloid cascade hypothesis has strongly influenced the view of sporadic AD progression, due to the observation that late-stage pathology is identical across familial and sporadic AD^8^. Following this view, the “ATN” framework has been recently proposed, aiming to shift AD from a symptom-based to a biomarker-based diagnostic entity. This research framework codifies AD progression via a specific set of biomarkers than can measure AD-related pathologies in vivo^9,10^. The set consists of amyloid-based markers (e.g. amyloid PET or CSF-based Aβ; the ‘A’ component), tau-based markers (e.g. tau PET or CSF-based tau; ‘T’) and markers of neurodegeneration or neuronal injury (e.g. MRI or FDG-PET; ‘N’).

Consistent with the amyloid cascade view, the ATN framework requires the presence of amyloid pathology for an individual to enter the AD continuum and therefore progress to subsequent stages of the disease. Any other pattern of biomarker abnormalities is incompatible with its view. However, both neuropathologic and in vivo studies have challenged the notion that amyloid pathology precedes tau pathology within AD. Neuropathology studies show that tau pathology (localized primarily within the entorhinal cortex) is present in roughly thirty percent of older subjects with no amyloid pathology^11^. Similarly, a recent amyloid and tau PET based biomarker study found that 45% of non-demented subjects were tau positive in entorhinal/hippocampal regions while being amyloid negative^12^. As a consequence, there is ongoing debate around what this focalized tau-based abnormality represents. While some suggest it relates to normal aging, a condition termed primary age-related tauopathy (PART^13^), or due to another proteinopathy (e.g. TDP-43), there is a possibility that this focalized tau abnormality consists of an alternative, ‘tau-first’ pathway towards AD dementia^14,15^. In fact, neuropathological studies have noted that the tau-based NFTs observed in PART are biochemically identical to those in AD^16,17^.

Taken together, these prior studies suggest that there may be two basic subtypes of pathology progression within AD: an amyloid-first subtype, embodying the amyloid cascade in which subjects develop Aβ plaques prior to NFTs, and a tau-first subtype, in which focalized NFT pathology precedes the appearance of plaques. To date, however, no study has identified and characterized these patterns across both post mortem and in vivo measures with the aim of understanding how these possible pathological pathways may be related to distinct concurrent cognitive and CSF-based abnormalities. Here, we address this knowledge gap using in vivo PET, CSF and cognitive measures from the Alzheimer’s Disease Neuroimaging Initiative (ADNI) and postmortem neuropathologic measures from the Religious Orders Study and Rush Memory and Aging Project studies (ROSMAP). We follow a data-driven paradigm to identify disease subtypes using the SuStaIn (Subtype and Stage Inference) algorithm^18^, which identifies groups of subjects with common patterns of disease progression from multi-modal cross-sectional data. SuStaIn is thus well-suited to inferring disease subtypes from both post mortem and in vivo measures. We consistently identify both amyloid-first and tau-first subtypes, supporting the dual-pathway hypothesis of AD progression. We find higher soluble TREM2 (sTREM2) measures in tau-first subjects’ CSF, suggesting increased microglial activation within this subtype. Our findings add to the understanding of AD progression and offer a precision tool for screening and stratifying patients in clinical settings.

## Results

### Tau-first subtype identified using post mortem measures

The ROSMAP study is an ongoing observational study of older adults that have agreed to annual clinical evaluation and cognitive testing as well as brain donation after death. Through December 31^st^, 2017 there were 3,414 subjects enrolled, with 1,717 deaths and 1,506 brain autopsies. We applied SuStaIn^18^, an unsupervised machine learning algorithm, to 1,211 ROSMAP study subjects’ post mortem immunohistochemistry measures of amyloid and tau pathologies in the eight available brain regions (hippocampus, entorhinal cortex, midfrontal cortex, inferior temporal, angular gyrus, calcarine cortex, anterior cingulate cortex, superior frontal cortex). These subjects had a complete set of measurements in all regions as well as a neuropathologically confirmed diagnosis of either normal aging or AD and an ante-mortem exclusion of other dementias.

As a preliminary step we estimated the number of subtypes that best explain the available observations. We allowed SuStaIn to infer one, two or three-subtype models and in each case evaluated the out-of-sample log likelihoods of held-out samples under ten-fold cross-validation. The two-subtype model improved upon the one-subtype model (14% median improvement, p = 1.95 × 10^-3^ via Wilcoxon signed-rank test) and there was a marginal further improvement with a the three-subtype model (3%; Figure 1a). Thus, we used the two-subtype model in subsequent analyses on the ROSMAP data.

**Figure 1.**
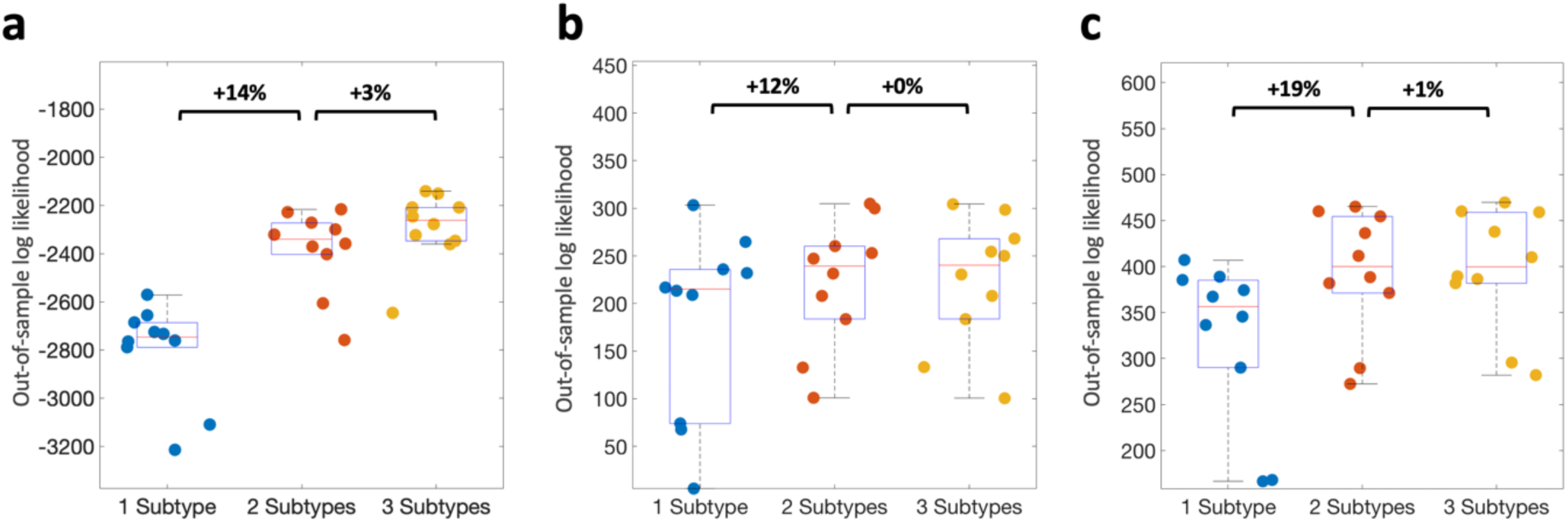
Out-of-sample data likelihood for each of ten cross-validation folds for one, two and three subtype models for **a** ROSMAP neuropathology-based analysis; **b** ADNI all-PET-based analysis; and **c** ADNI PET- and-CSF-based analysis.

Figure 2 depicts positional variance diagrams (PVDs) estimated by SuStaIn for the two subtypes identified in ROSMAP. PVDs visualize event sequence uncertainties as a matrix; each row is a histogram of the fraction of times an event occurs at a given position across a set of Markov-chain Monte Carlo (MCMC) event-sequence samples. Here we have 16 events (amyloid and tau across eight brain regions) and hence as many distinct stages of progression within each subtype. Within the two-subtype model, cognitively normal subjects had a median stage of four in the first subtype and three in the second; AD subjects had a median stage of 11 in the first subtype and 12 in the second (Supplementary Figure 2a, 2d).

**Figure 2.**
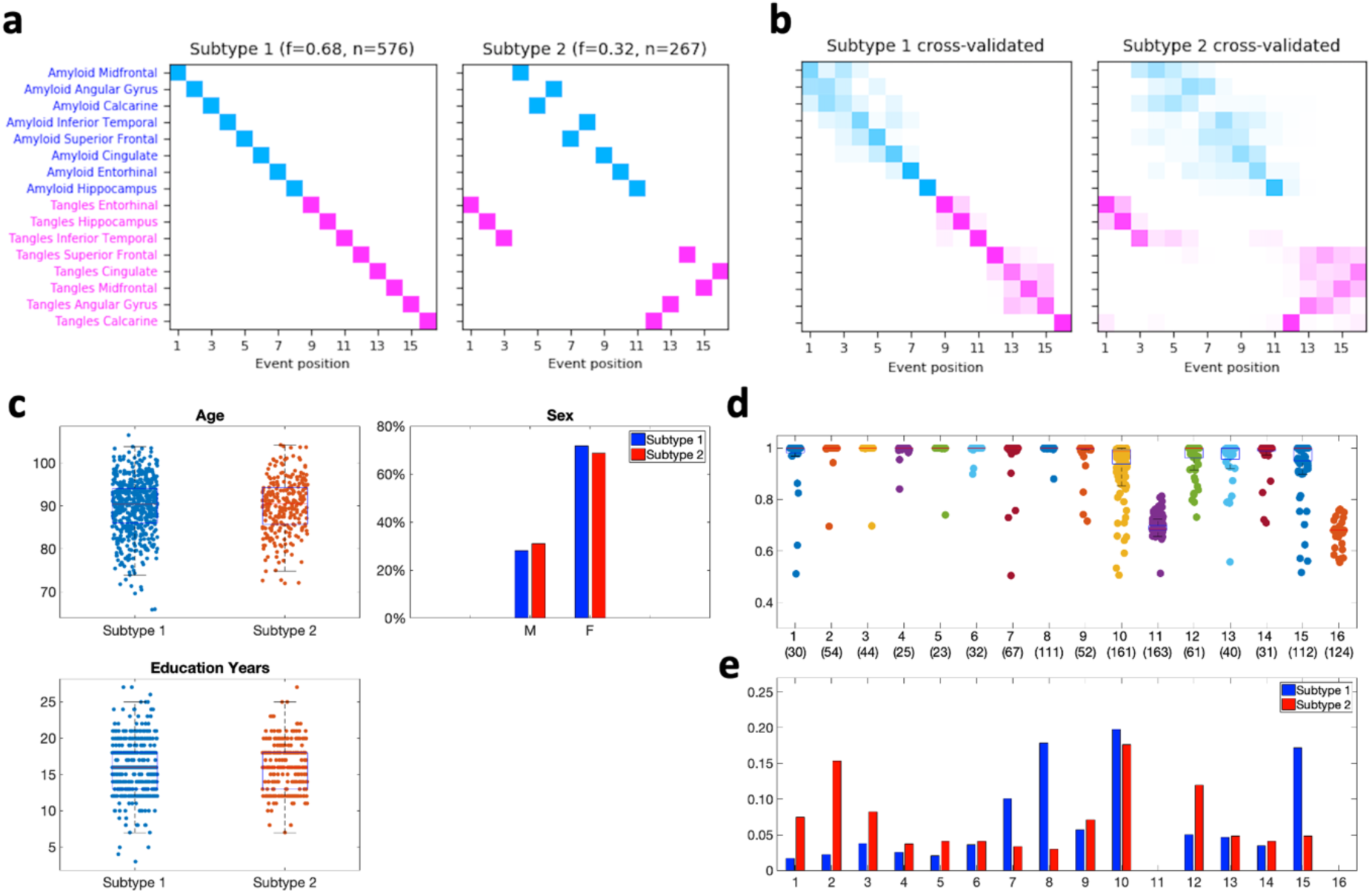
Inferred subtypes and stages in the ROSMAP neuropathology-based analysis. **a** positional variance diagrams (PVDs) for two-subtype Sustain model of regional amyloid beta and tau-based neurofibrillary tangle pathology spread; **b** PVDs from ten fold cross-validated version of the same model, providing an estimate of event uncertainty within both subtypes; c demographic breakdown of subtypes by age, percentage of each sex and education years; **d** boxplots of subtype probability at each stage, with number of subjects at each stage shown in parentheses on the x-axis; and e histograms of fraction of subjects at each stage per subtype with the 163 subjects at stage 11 and the 124 subjects at stage 16 removed due to high uncertainty in their most likely subtype (see d).

We estimated the prevalence of each subtype using subjects most representative of their subtype. As SuStaIn is a probabilistic algorithm that assigns a non-zero probability of belonging to each subtype and stage for every subject we chose to retain only those with a relatively high probability of most likely subtype. In general, these are subjects who are in middle stages of progression; those at the extremes have either no pathology (very early stages) or abnormality across all markers (very late stages) and thus the notion of being in a particular subtype is not meaningful for them. We therefore removed 81 subjects at stage zero and 124 subjects at stage 16. We also removed 163 subjects at stage 11 as the model indicates that at these stages the two subtypes are also not distinguishable (Figure 2d). Thus, we retained 843 out of 1,211 subjects for further analysis of subtypes.

SuStaIn found an amyloid-first subtype (68%, 576 of 843 subjects) in which amyloid pathology begins in the midfrontal, angular gyrus and calcarine regions of the cortex. Once amyloid has spread throughout the cortex, tau pathology manifests, beginning in the entorhinal, hippocampus and inferior temporal regions and then spreading throughout the remaining cortex. The second identified subtype is a tau-first subtype (32%, 267 of 843 subjects) in which neurofibrillary tau tangle pathology similarly begins in the entorhinal, hippocampus and inferior temporal regions and is followed by a sequence of amyloid pathology progression that closely resembles that of the amyloid-first subtype. After the amyloid stages, tangle formation then expands to the remaining cortex (calcarine, angular gyrus, superior frontal, midfrontal, and cingulate regions; Figure 2a).

We used cross-validation to get a more robust estimate of event sequence uncertainties as MCMC tends to underestimate the uncertainty of events within each subtype^19^. We find similar subtypes with greater uncertainty in the ordering of amyloid pathology in both subtypes and higher uncertainty in late-stage tau pathology in the second subtype (Figure 2b).

### No differences in demographic factors or comorbid pathologies between subtypes

We investigated basic demographic differences between subtypes (depicted in Figure 2c) while controlling for the effect of subjects’ stage. We found no statistical differences related to age at death, years of education or sex between subjects in subtype one (amyloid-first) compared to subtype two (tau-first). We also tested for differences in comorbid pathologies, finding no differences in TDP-43, cerebral amyloid angiopathy (CAA), cerebral atherosclerosis or arteriolosclerosis and no difference in the presence of hippocampal sclerosis (Supplementary Table 3).

### In vivo PET and CSF based subtyping models are consistent with neuropathology

We used in vivo measures from the Alzheimer’s Disease Neuroimaging Initiative (ADNI), a long-running observational study tracking healthy, cognitively impaired and AD-diagnosed older adults. We performed two separate SuStaIn-based analyses using ADNI study measures. The first (the ‘all-PET-based analysis’) was an in vivo analog to the ROSMAP neuropathologic analysis. We used all subjects who had concurrent amyloid PET and tau PET images (n = 445), showing that, as in the ROSMAP analysis, a two-subtype model of pathology spread improved upon a one-subtype model (12% median improvement in out-of-sample log likelihood, p = 3.91 × 10^-3^ via Wilcoxon signed-rank test) and there was no further improvement with a three-subtype mode (0%; Figure 1b). The in vivo data therefore confirmed the appropriateness of a two-subtype model. Cognitively normal subjects had a median stage of zero (out of a total of 21 stages; see Figure 3) in the first subtype and were not staged in the second, MCI subjects had a median stage of one in the first subtype and two in the second and AD subjects had a median stage of 14 in the first subtype and 15 in the second (Supplementary Figure 2b, 2e).

**Figure 3.**
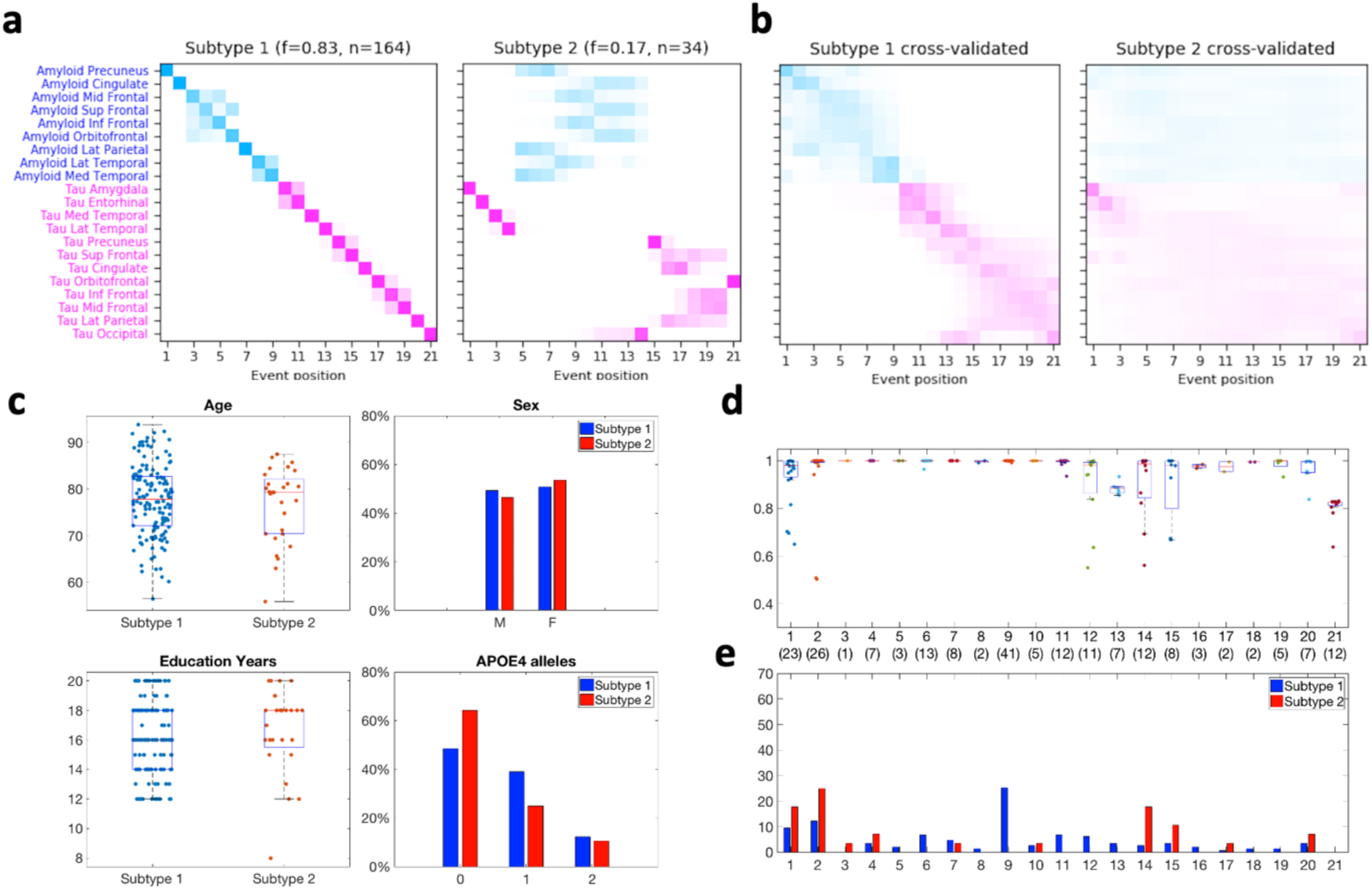
Inferred subtypes and stages in the ADNI all-PET-based analysis. **a** positional variance diagrams (PVDs) for two-subtype SuStaln model, showing the most likelihood sequence of amyloid PET and tau PET based pathology spread; **b** PVDs from ten fold cross-validated version of the same model, providing an estimate of event uncertainty within both subtypes; **c** demographic breakdown of subtypes by age, percentage of each sex, education years and percentage of APOE4 alleles; **d** boxplots of maximum likelihood subtype probability at each stage, with number of subjects at each stage shown in parentheses on the x-axis; and e histograms of fraction of subjects at each stage per subtype with 12 subjects at stage 21 removed due to high uncertainty in their most likely subtype (see d).

As in the neuropathologic analysis, we estimated the prevalence of each subtype by first removing 235 subjects at stage zero and 12 subjects at stage 21, leaving 198 subjects. We found an amyloid-first subtype (83%, 164 of 198 subjects) in which amyloid pathology begins in the precuneus, cingulate, and frontal (inferior, middle, superior) cortical regions, and a tau-first subtype (17%, 34 of 198 subjects) in which tau pathology begins in the amygdala, entorhinal, and temporal (medial and lateral) regions (Figure 3a). As in the neuropathologic analysis, in both in vivo subtypes we found that tau spreads beyond the hippocampus, amygdala and temporal lobe only after amyloid has spread throughout the cortex. Under cross-validation we inferred similar progression patterns: an amyloid-first subtype and a tau-first subtype with greater uncertainty in the event orderings within each subtype, particularly within the tau-first subtype (Figure 3b).

The small number of subjects falling into the tau-first subtype (n = 34) limits the ability to precisely estimate the event ordering within this subtype. This also limited our ability to detect demographic, cognitive and CSF-based differences between subtypes. Therefore, our second ADNI-based analysis (‘PET-and-CSF-based analysis’) addressed this by increasing the number of subjects using CSF-based regional tau prediction as a proxy for tau PET. In this case we used the available 999 subjects who had both amyloid PET and CSF-based (total and phosphorylated) tau measures at a single visit. We selected the two-subtype model as in the previous analyses (Figure 1c). Cognitively normal subjects had a median stage of zero in the first subtype and stage three in the second, MCI subjects had a median stage of four in first subtype and three in the second and AD subjects had a median stage of 12 in the first subtype and four in the second (Supplementary Figure 2c, 2f).

To estimate the prevalence of each subtype, we excluded 459 subjects at stage zero and 279 subjects at stage 12 (Figure 4d), leaving 261 subjects for further analysis. We similarly found an amyloid-first subtype (62%, 162 of 261 subjects) in which amyloid pathology (measured via amyloid PET) begins in the frontal (inferior, middle, superior), cingulate and precuneus regions, and a tau-first subtype (38%, 99 of 261 subjects) in which tau pathology (estimated from CSF and demographic information) begins in Braak I/II, III/IV and V/VI regions (Figure 4a, 4b).

**Figure 4.**
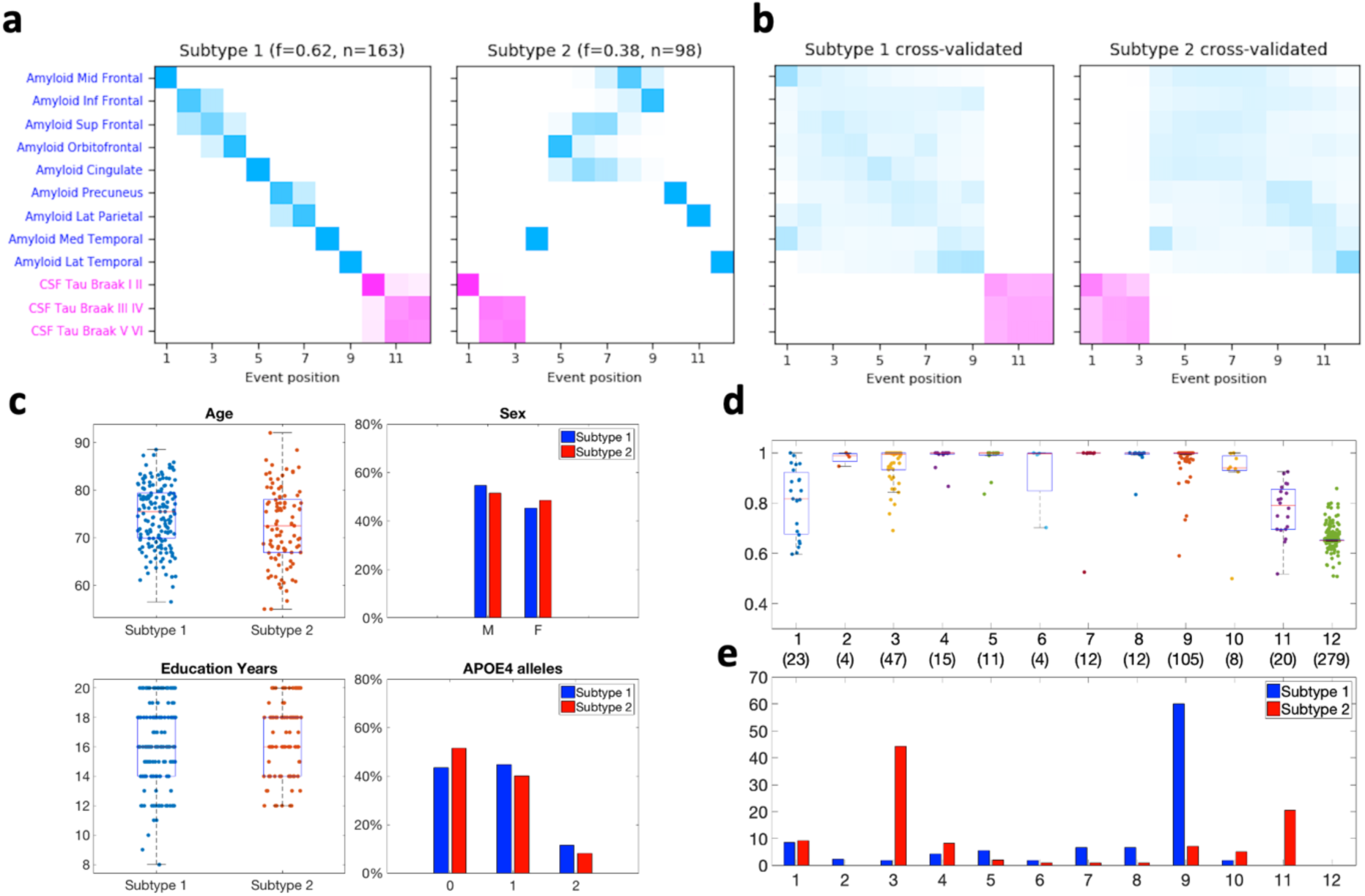
Inferred subtypes and stages in theADNI PET-and-CSF-based analysis. **a** positional variance diagrams (PVDs) for two-subtype SuStaln model, showing the most likelihood sequence of amyloid PET based and CSF tau based pathology spread; **b** PVDs from ten fold cross-validated version of the same model, providing an estimate of event uncertainty within both subtypes; **c** demographic breakdown of subtypes by age, percentage of each sex, education years and percentage of APOE4 alleles; and **d** boxplots of subtype probability at each stage, with number of subjects at each stage shown in parentheses on the x-axis; and **e** histograms of fraction of subjects at each stage per subtype with 279 subjects at stage 12 removed due to high uncertainty in their most likely subtype (see d).

### CSF based tau, amyloid beta and sTREM2 differences between subtypes

We leveraged the deep phenotyping data available in ADNI and performed logistic regression-based analyses to test for group differences between our data-driven in vivo subtypes. For this we used the PET-and-CSF-based analysis due to its higher sample size and overall similarity to the PET-based analysis cohort along most demographic and diagnostic measures (Supplementary Table 2). There was one important difference between cohorts: subjects in the all-PET-based analyses were on average 3.3 years older, 95% CI = [1.9, 4.6], p = 3.42 × 10^-6^, and measured 3.6 years further out from baseline, 95% CI = [3.2, 4.1], p < 1 × 10^-6^, than those in the PET-and-CSF-based analysis. This is due to tau PET being introduced in a later stage of the ADNI study (ADNI-3^20^).

In the first subtype comparison, which included 259 of 261 subjects who had both CSF-based tau and amyloid as well as cognition, we found a small difference in age between subtypes (tau-first subjects 2.3 years younger, 95% CI = [0.5, 4.1], odds ratio 0.57, p = 0.02; Figure 4c), no differences in memory or executive function (Supplementary Table 4; Figure 5a) but found, as anticipated, substantial differences in tau (tau-first subjects 115.1 pg/mL higher, i.e. more abnormal, 95% CI = [94.0, 136.1], odds ratio 7.18, p < 1 × 10^-6^) and amyloid beta (tau-first subjects 689.8 pg/mL higher, i.e. less abnormal, 95% CI = [545.5, 834.2], odds ratio 2.83, p = 7.53 × 10^-4^; Figure 5b).

**Figure 5.**
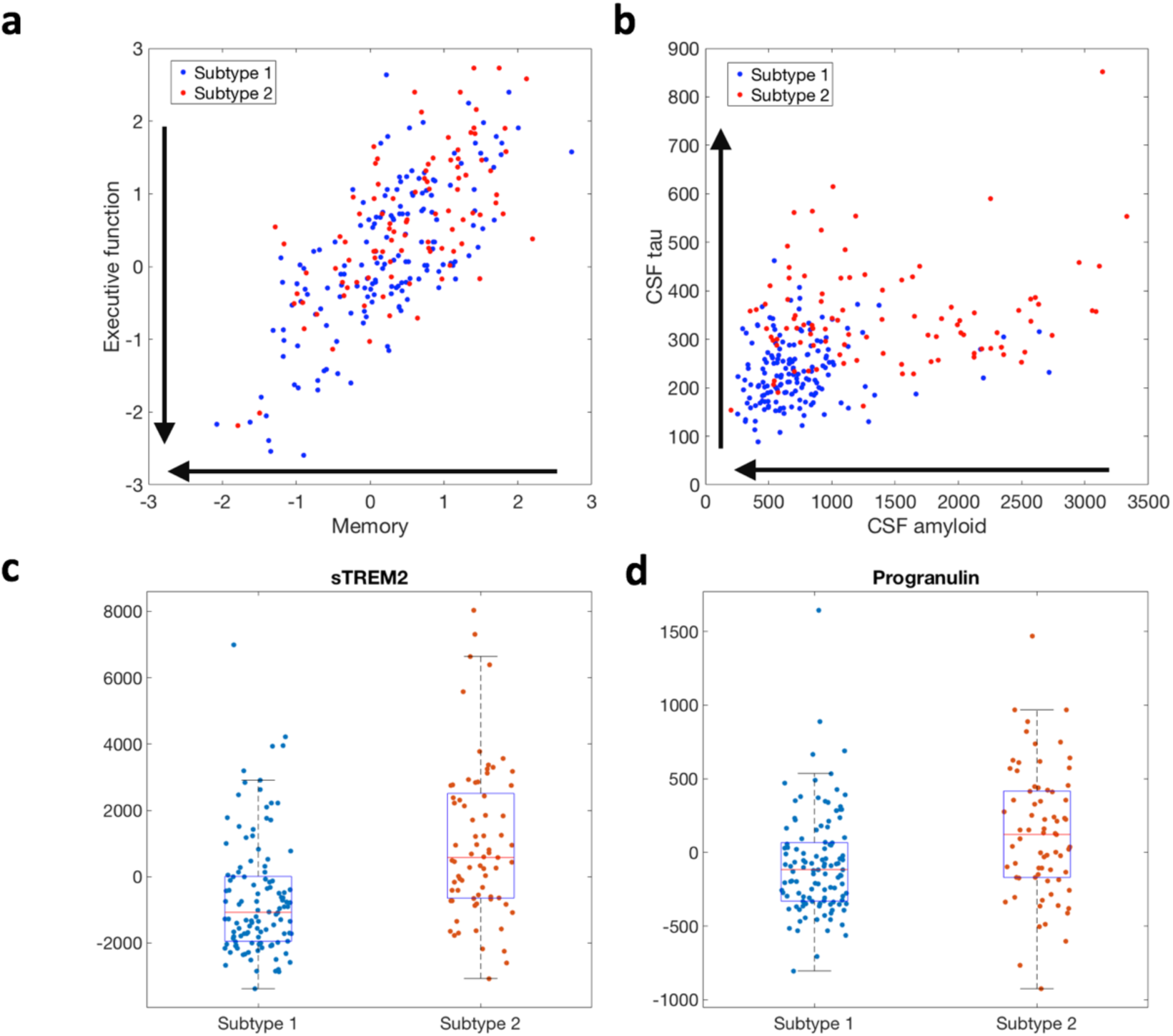
Subtype differences in theADNI PET-and-CSF-based analysis. **a** scatter plot of memory and executive function measures for subtype one (amyloid-first) and subtype two (tau-first) subjects, showing no apparent differences in cognition between subtypes; **b** scatter plots for CSF amyloid and tau measures, showing that subtype one (amyloid-first) subjects’ CSF amyloid measures are generally lower (more abnormal) while subtype two (tau-first) subjects’ CSF tau measures are generally higher (more abnormal); **c** boxplots of differences in sTREM2 across subtypes and **d** same for progranulin. Arrows in **a, b** show direction of increasing abnormality for each measure.

**Figure 6.**
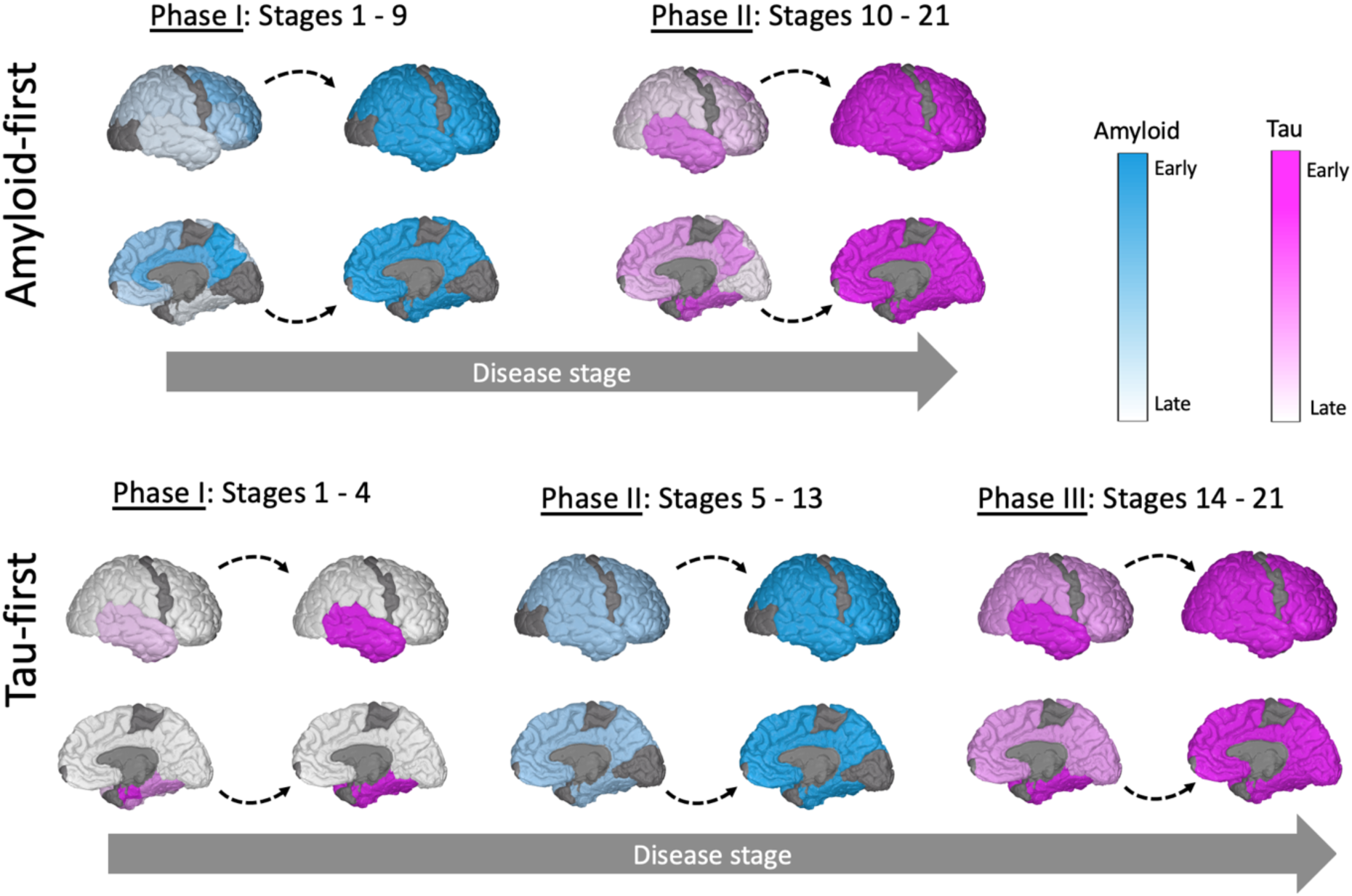
Phases of amyloid and tau pathology spread within the two identified subtypes in the ADNI all-PET-based analysis, depicted in Figure 3a. Within the amyloid-first subtype, amyloid pathology spreads throughout the cortex beginning in the precuneus, cingulate and frontal lobe (phase I, top). Tau then spreads from the amygdala (not shown), entorhinal and medial and lateral temporal lobe to remaining cortex (phase II, top). Within the tau-first subtype, tau pathology in the amygdala, entorhinal cortex, and medial and lateral temporal lobe precedes amyloid pathology (phase I, bottom). Amyloid subsequently spreads throughout the cortex (shown in an intermediate blue color due to high uncertainty in the pattern of progression; phase II, bottom). Tau then spreads throughout the remaining cortical regions (shown in intermediate magenta due to high uncertainty; phase Ill, bottom). Regions in grey are not included in the model. Drawn using BrainPainter.

In the second subtype comparison, involving a subset of 190 of 261 subjects with CSF based sTREM2 and progranulin, we found a small difference in age (tau-first subjects 3.0 years younger, 95% CI = [-5.1, -1.0], odds ratio 0.58, p = 2.83 × 10^-3^) as well as a difference in sTREM2 (tau-first subjects 1652.8 pg/mL higher, i.e. more abnormal, 95% CI = [1069.3, 2236.3], odds ratio 4.18, p = 2.92 × 10^-5^; Figure 5c) but no difference in progranulin (Supplementary Table 5; Figure 5d).

Across both comparison analyses we found no statistical differences in sex, number of APOE4 alleles, years of education, nor model stage (Figure 4c, 4e).

## Discussion

While amyloid- and tau-based pathologies have long been established as the main pathological hallmarks of AD, the heterogeneity within the spatiotemporal progression of these pathologies has yet to be fully characterized. Here, for the first time, we used a fully data-driven model on two large cohorts with complementary in vivo and post mortem measures, to show the presence of two subtypes of amyloid and tau pathology progression within sporadic AD. We consistently found ‘amyloid-first’ and ‘tau-first’ subtypes (Figure 1). In the ‘amyloid-first’ subtype, extensive cortical amyloid pathology precedes tau pathology. In the ‘tau-first’ subtype, localized tau pathology precedes the spread of amyloid. In both datasets, the majority of subjects aligned with the amyloid-first subtype (68% of subjects based on immunohistochemistry, 83% based on PET, 62% based on CSF and PET) while a non-negligible minority of subjects fell into the tau-first subtype (32% immunohistochemistry, 17% PET, 38% CSF and PET). Importantly, we found that early tau pathology in the tau-first subtype is localized within the temporal lobe, hippocampus and amygdala (Figures 2a, 2b, 3a, 3b) and that in both subtypes amyloid pathology spreads throughout the cortex before tau pathology spreads beyond the temporal lobe to other parts of the cortex (Figure 621). This finding is consistent with neuropathologic studies showing that while the initial formation of tau and amyloid pathologies may be independent of each other, widespread amyloid pathology precedes and likely facilitates the spread of tau pathology beyond the medial temporal lobe (MTL) and limbic areas^11,22^.

Previous studies of the spread of amyloid pathology in AD have used both post mortem and in vivo measures. Braak and Braak, using silver staining of post mortem brains, observed that amyloid pathology is first found in the basal frontal, temporal and occipital lobes before spreading throughout the cortex, with sensorimotor regions and subcortical structures such as the hippocampus becoming gradually involved^23^. Thal et al., using both silver staining and immunohistochemistry, found a similar pattern^24^. Fantoni et al. summarized a number of recent in vivo amyloid PET based studies, noting that the first regions with detectable abnormality are the medial frontal and cingulate regions, followed by a general spreading throughout the cortex, the striatum, hippocampus, other subcortical regions and ultimately the cerebellum^25,26,27,28,29,30,31,32^. The pattern of amyloid accumulation that we inferred was consistent with these studies. In our analysis of post mortem immunohistochemistry data, we found two data-driven subtypes, each having similar patterns of amyloid spread. The earliest abnormalities were in the midfrontal cortex followed by the angular gyrus, calcarine, inferior temporal, superior frontal and other cortical regions, with hippocampus as the latest event. Cross-validation suggests less certainty in the ordering (Figures 2a, 2b). In our analysis of in vivo amyloid PET data, we inferred a pattern of spread from the precuneus and cingulate to the frontal, parietal and temporal lobes of the cortex, with cross-validation again suggesting less certainty in the ordering of the sequence (Figures 3a, 3b). Our inferences of the ordering of in vivo amyloid progression within the tau-first subtype were limited by the number of subjects in this group.

Previous studies have similarly used both post mortem and in vivo measures to characterize the spread of tau pathology in AD. Braak and Braak developed the standard six-stage system used to describe NFT spread based on neuropathologic examination: at stages I and II NFTs initially form within the transentorhinal cortex, at stages III and IV NFTs spread to the entorhinal cortex, hippocampus and amygdala and by stages V and VI pathology has spread throughout the frontal, parietal and occipital cortex, mostly sparing the sensorimotor region^23^. Several studies have shown that these Braak stages are detectable in vivo using tau PET^33,34,27^. In our analysis of post mortem immunohistochemistry data, we found NFT pathology beginning in the entorhinal, hippocampus and inferior temporal regions (corresponding to Braak I-III/IV regions) before spreading throughout the remaining cortex (Braak IV-VI regions). While both of our data-driven post mortem subtypes broadly conformed to this pattern, our results suggest that the tau-first subtype deviates from the expected Braak progression after the initial accumulation of tau and the spread of amyloid. Within this subtype NFTs unexpectedly appear in the calcarine cortex (within the occipital lobe, Braak stage VI) prior to? NFTs in the frontal, parietal and cingulate regions, corresponding to stages IV and V (Figures 2a, 2b). Our analysis of in vivo PET data was in concordance with our analysis of post mortem data: amygdala, entorhinal and both medial and lateral temporal regions appear early, followed by other cortical areas. There is a similar subtype-specific pattern: tau spreading within the amyloid-first subtype closely conforms to Braak staging, while tau pathology in the tau-first subtype deviates from Braak staging, with pathology in the occipital lobe appearing prior to frontal, parietal, and cingulate regions (Figures 3a, 3b). Together, these findings point to a subtype-specific pattern of tau spread throughout the cortex, with variability in later-stage progression.

When using CSF-based predictions of regional tau pathology, we found that late-stage tau pathology (within Braak V/VI regions) precedes amyloid pathology in the tau-first subtype, which is inconsistent with our post mortem and PET models (compare Figures 4a, 4b with Figures 2 and 3). Mattson et al.^35^ noted that CSF-based tau measures become abnormal during preclinical AD stages but do not track AD progression into later stages, while tau PET based measures are more closely associated with the neurodegeneration and cognitive decline seen in clinical AD stages. This is consistent with our observation that the regression models trained to predict regional tau PET SUVRs from CSF-based tau and demographics tend to plateau at higher values and have difficulty resolving between middle and late stage Braak pathology (see Supplementary Figure 1). Comparing this model to the immunohistochemistry and PET-only models, we see that this limited the ability of the CSF-based model to detect late-stage tau pathology in the tau-first subtype.

One of the strengths of our study was in the use of complementary information from post mortem and in vivo measures to infer subtypes. We found robust amyloid-first and tau-first subtypes across these measures which enabled, for the first time, comparison of subtypes across a variety of demographic, cognitive, and comorbid pathology measures while control for subjects’ stage of progression within their respective subtype. We found no differences between subtypes in demographic factors or genetic risk (sex, years of education, APOE4 alleles) and only a small difference in age (amyloid-first subjects several years older on average) in the PET-and-CSF-based model (Supplementary Tables 3, 4, and 5). We also found no differences in comorbid pathologies related to cerebrovascular function, TDP-43 and hippocampal sclerosis, implying that these pathologies may develop in similar ways in both subtypes (Supplementary Table 3).

Our findings suggest that a substantial number of “PART-like” subjects are actually in the early stages of the tau-first AD subtype that we have identified. We found no statistical differences in memory or executive function between our amyloid-first and tau-first subtypes, implying that tau-first subjects develop cognitive impairment as they progress to later stages just as amyloid-first subjects do (Figure 5a, Supplementary Table 4). Similarly, a recent PET based study found that tau-first subjects show signs of early cognitive impairment, supporting the notion that PART (i.e. MTL-localized tau pathology with no detectable amyloid pathology) is within the AD continuum. Further to this, our post mortem immunohistochemistry model shows that while many tau-first subjects may not progress beyond localized tau pathology (i.e. up to stage four within their subtype), others will go on to develop amyloid pathology (i.e. stages five and above; Figures 2a, 2e, 3a, 3e). Future work includes understanding how amyloid-first and tau-first subtypes are related to the three subtypes of AD-related neurodegeneration that have been consistently identified across post mortem and in vivo studies^36,37,38,18^.

The amyloid-cascade hypothesis maintains that amyloid pathology, driven by cellular processes related to lipid metabolism, microglial activation and endocytosis, initiates downstream tau and neurodegeneration. In contrast, the dual-pathway hypothesis proposes instead that these cellular processes may drive amyloid and tau pathologies along independent pathways^14,15^. Our data-driven models, based on both in vivo and post mortem measures, lend support to the dual-pathway hypothesis (Figure 7). We note however, that amyloid pathology may have a similar catalyzing role in both subtypes: facilitating the spread of tau pathology beyond the MTL and limbic areas. Our finding that CSF-based soluble TREM2, thought to be a maker of microglia function, is increased in tau-first subjects (consistent with previous findings^39^), provides a first step in directly relating these subtypes to upstream cellular processes (Supplementary Table 5). Further work is needed using measures of lipid metabolism, microglia and endocytosis as well as genome-wide association studies (GWAS) to better understand how these subtypes differ in terms of cellular processes and their upstream genetic risk factors.

**Figure 7.**
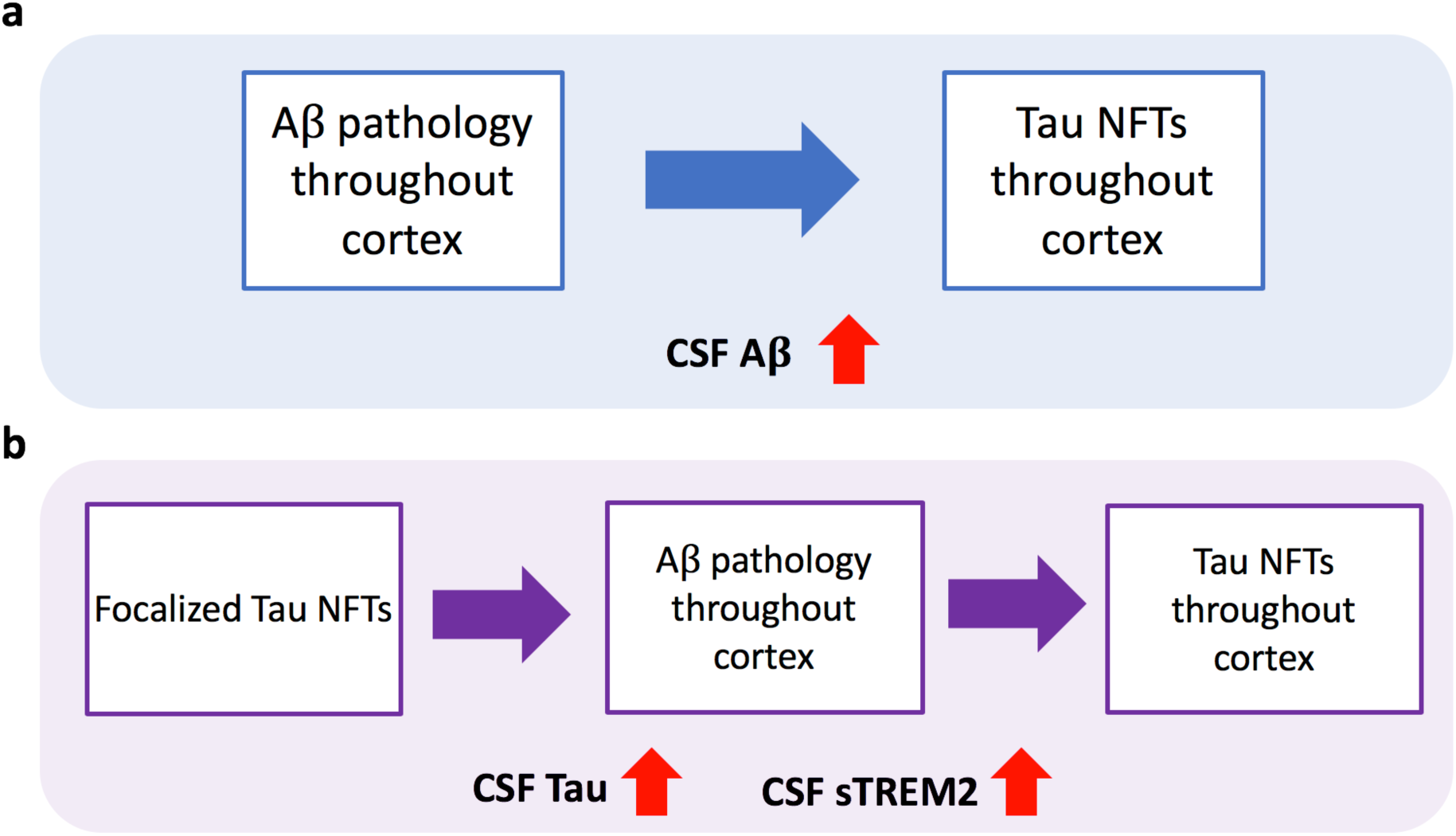
Characterization of identified subtypes. Data-driven models consistently identify two subtypes of amyloid and tau pathology progression across both post mortem and in vivo measures. This first, depicted in panel **a,** is an amyloid-first subtype, in which Aβ-based pathology spreads throughout the cortex prior to the spread of tau-based neurofibrillary tangles (NFTs). Within this subtype CSF-based Aβ is more abnormal relative to tau-first subtype subjects. The second, depicted in panel **b,** is a tau-first subtype in which focalized tau-based NFTs within Braak I-III/IV regions precede the spread of Aβ pathology throughout the cortex, which is followed by the spread of tau-based NFTs into the remaining cortex (Braak IV-VI regions). CSF-based tau and sTREM2 are more abnormal within this subtype relative to those in the amyloid-first subtype.

In summary, we have identified a tau-first subtype of sporadic AD progression using data-driven disease subtyping models trained on both in vivo and post mortem measures. These findings have important implications for interventional trials targeting amyloid-β and tau pathologies.

## Methods

### ADNI dataset

Data used in the preparation of this article were obtained from the Alzheimer’s Disease Neuroimaging Initiative (ADNI) database (adni.loni.usc.edu). The ADNI was launched in 2003 as a public-private partnership, led by Principal Investigator Michael W. Weiner, MD. The primary goal of ADNI has been to test whether serial magnetic resonance imaging (MRI), positron emission tomography (PET), other biological markers, and clinical and neuropsychological assessment can be combined to measure the progression of mild cognitive impairment (MCI) and early Alzheimer’s disease (AD). For up-to-date information, see www.adni-info.org.

We downloaded regional amyloid PET (AV-45) standardized update value ratios (SUVRs), partial volume corrected regional tau PET (AV-1451) SUVRs and cerebrospinal fluid (CSF) based measures of amyloid (Aβ1–42), total tau and phosphorylated tau (pTau) from the ADNI database^40,41^. We also downloaded the ADNIMERGE table, containing demographic information (age, sex, years of education, number of APOE4 alleles), and diagnostic labels (cognitively normal/MCI/AD). We downloaded composite measures of memory (ADNI-MEM^42^) and executive function (ADNI-EF^43^). Finally, we downloaded CSF based measures of sTREM2 and progranulin, using the measurements made via the MSD ELISA platform that were subsequently corrected by plate-specific factors^44^. The ADNI database was last accessed on February 6^th^, 2020.

### ROSMAP dataset

We used post mortem neuropathology data from the Religious Orders Study (ROS) and Rush Memory and Aging Project (MAP) studies, collectively referred to as ROSMAP, which we obtained from the Rush Alzheimer’s Disease Center (RADC)^45^. We used molecularly-specific immunohistochemistry based measures of amyloid beta protein and neuronal neurofibrillary tangles (associated with abnormally phosphorylated tau protein) both measured in eight brain regions (hippocampus, entorhinal cortex, midfrontal cortex, inferior temporal cortex, angular gyrus, calcarine cortex, anterior cingulate cortex and superior frontal cortex) along with demographic information (age at death, sex, education years), final (in vivo) clinical diagnosis of AD (NINCDS-ARDRA^46^) and (post mortem) neuropathologic diagnosis of AD (NIA-Reagan Criteria^47^).

We also compared comorbid pathologies between subtypes, using RADC’s stagings of TDP-43, cerebral amyloid angiopathy (CAA), cerebral atherosclerosis and arteriolosclerosis based pathologies, each ranging zero (no pathology) to three (severe pathology), along with the presence or absence of hippocampal sclerosis.

### Disease progression modeling

We used Subtype and Stage Inference (SuStaIn), a probabilistic machine learning based method, to characterize the heterogeneity of amyloid and tau pathology progression in AD. SuStaIn infers multiple patterns of disease progression (i.e. subtypes) as well as individuals’ disease stages from cross-sectional data^18^. The SuStaIn model as introduced by Young et al.^18^ uses a data likelihood based on how far a biomarker measurement deviates from normality, with an associated set of z-score based events (e.g. one, two or three z-scores away from control population mean) for each biomarker. Note that in biomarkers where controls have very little abnormality (e.g. amyloid load in cognitively normal APOE4 negative subjects, as in our work), the resulting z-scores in patients can become large owing to the small amount of variance in the control population. In such cases it is more sensible to use a separate distribution to describe patients’ measurements and define an event as a biomarker going from normal to abnormal (as in the event-based model; EBM^48,19^). Formally, we use *P*(*x_ij_*|*E_i_*) and *P*(*x_ij_*|¬*E_i_*), the likelihoods of measurement *x_ij_* of subject *j* in the case where biomarker *i* has or has not become abnormal (event *i* has occurred, *E_i_*, or event *i* has not occurred, ¬*E_i_*, respectively) within the EBM-based data likelihood,

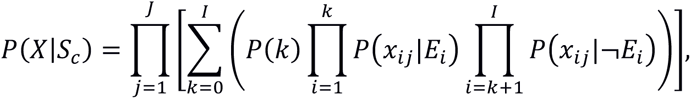

where *S*_#_ is the sequence of events within a particular subtype *c*, *X* = {*x_ij_*|*i* = 1, …, *I*; *j* = 1, …, *J*} is the set of measurements over *I* biomarkers and *J* subjects and *P*(*k*) is the prior likelihood of being at stage *k*, assumed to be uniform. The remainder of SuStaIn’s model fitting procedure is as described in detail in Young et al.^18^ It consists of an iterative procedure that simultaneously optimizes subtype event sequences (finding event sequence *S*_#_ for each subtype *c*) and subtype membership (assigning a probability of being in subtype *c* to every subject) for a pre-selected number of subtypes *C*. For each subtype, a Markov chain Monte Carlo (MCMC) based procedure is used to estimate the uncertainties of event orderings, which can be visualized with a positional variance diagram (PVD)^48^. In our work we fit *P*(*x_ij_*|*E_i_*) and *P*(*x_ij_*|¬*E_i_*) for each biomarker using the kernel density estimation based mixture modeling approach described by Firth et al.^49^

We optimized the number of subtypes (*C*) in an iterative manner using ten-fold cross-validation. For each fold we ran SuStaIn on the training data and evaluated the average out-of-sample log likelihood of the held-out testing data, so that we had ten measures for each model. We could thus compare a *C*-subtype model to a (*C* + 1)-subtype model via the median relative improvement in log likelihood across folds. We stopped iterating *C* when the (*C* + 1)-subtype model no longer substantially improved upon the *C*-subtype model, making the *C*-subtype model the more parsimonious choice.

We used the same ten-fold cross-validation procedure to generate cross-validated PVDs in order to visualize the uncertainty in the inferred ordering of events within a subtype. To do this, we compared each fold’s inferred subtypes to an overall model built on all samples, finding the best one-to-one matching between the folds’ inferred sequences and the overall model’s using Kendall’s tau as a similarity metric. Based on this mapping we concatenated matching subtypes’ MCMC samples across folds and calculated PVDs. The code for this procedure is available in our implementation of SuStaIn (see Data Availability section).

### ROSMAP based analysis

We performed a SuStaIn-based analysis using ROSMAP’s amyloid beta and neurofibrillary tangle measures in eight brain regions. Out of a total of 1,338 subjects who had complete set of measures in all regions we retained 1,226 subjects with a last clinical diagnosis of no cognitive impairment (CI) or either MCI or AD with no other cause of CI. The excluded 112 subjects had other dementia or cause of CI. We used subjects’ neuropathologic diagnosis for the remaining analysis, defining controls as those with low likelihood or no AD by NIA-Reagan pathology criteria and AD subjects as those with intermediate or high likelihood of AD using the same criteria.

We took the square root of each measure to improve normality and then corrected each measure for the effect of normal aging and demographics by training a regression model on control subjects’ values against age at death, sex and education years. We then regressed out the effect of normal aging from both controls and AD subjects, excluding 15 subjects who had at least one regional measure that was at least four standard deviations from the control or AD distribution mean. We then performed both mixture modeling and SuStaIn modeling using 1,211 subjects (418 controls/793 AD).

We investigated subtype differences using logistic regression with subtype (coded as zero for amyloid-first subtype, one for tau-first) as the dependent variable and demographic factors (age at death, sex, education years) and comorbidities (stages of TDP-43, cerebral amyloid angiopathy (CAA), cerebral atherosclerosis and arteriolosclerosis pathologies as well as presence of hippocampal sclerosis) as explanatory variables.

### ADNI based analyses

We performed two separate SuStaIn-based analyses using cross-sectional data from ADNI. The first was an all-PET-based analysis, in which we used nine regional amyloid PET (AV-45) SUVRs and twelve tau PET (AV-1451) SUVRs, many of which were volume-weighted combinations of several Freesurfer-based SUVRs (see Supplementary Table 1)^50,51^. We excluded the hippocampal tau PET SUVR as this region is suspected to be contaminated by off-target binding in the choroid plexus^52^. We reference normalized all SUVRs as recommended: for amyloid PET we used a composite reference region made up of the whole cerebellum, brainstem/pons and eroded subcortical white matter; for tau PET we used the inferior cerebellar grey matter^53,54^. We formed biomarkers for further analysis by log transforming these normalized SUVRs to improve normality.

For each biomarker we removed the effect of normal aging and demographic factors by training a regression model for each biomarker’s values against age, sex and education years in a control population of cognitively normal, APOE4 negative, global amyloid SUVR negative (whole cerebellum normalized summary SUVR < 1.11 cut-off) subjects. We then regressed out the signal due to these factors from all subjects’ measurements. Out of a total of 1515 subjects with amyloid PET or tau PET scans, we used 389 subjects to build mixture models of corrected amyloid PET biomarkers. These were 205 controls (as define above) and 184 AD subjects. We used 180 subjects for corrected tau PET mixture models: 116 CN, APOE4 negative, global amyloid SUVR negative subjects and 64 AD subjects. For SuStaIn modeling we used 445 subjects that had both amyloid and tau PET images at the same visit: 115 CN subjects, 290 with mild cognitive impairment (MCI) and 40 AD subjects. No outliers were removed in this analysis.

The second analysis was a PET-and-CSF-based analysis, substituting CSF-based prediction of tau PET SUVR in place of actual tau PET to give a larger dataset of 1001 subjects, all of whom had both CSF and amyloid PET at the same visit. The purpose of this analysis was to increase the sample size for comparing subtypes as many more subjects in ADNI have concurrent amyloid PET and CSF than concurrent amyloid and tau PET. Rather than using CSF total tau and pTau directly however, we approximated the first analysis by using the demographic and CSF measures to predict composite tau PET regions, training a regression-based prediction model using the 356 subjects with both tau PET and CSF measured at the same visit. We trained three separate linear regression models with composite SUVRs (regions related to I/II, III/IV or V/VI Braak stages) as the dependent variable and age, sex, education years, log CSF total tau and log CSF pTau as the explanatory variables. We used each model to predict tau SUVRs in the 645 subjects which did not have tau PET. Following this we formed a set of 12 biomarkers using the same nine amyloid PET SUVRs as before along with these three CSF-based tau markers, all of which were corrected for aging and demographics as before. We performed mixture modeling for each corrected biomarker using 193 CN, APOE4 negative, global amyloid SUVR negative subjects and 158 AD subjects. Prior to SuStaIn modeling we removed two outlying MCI subjects who had at least one biomarker that was at least four absolute standard deviations from either the control or AD distribution mean, so that we performed SuStaIn modeling with 999 subjects (366 CN/475 MCI/158 AD).

We investigated differences in demographic factors (age, sex, education years), CSF based amyloid beta and total tau, CSF-based soluble TREM2 (sTREM2) and progranulin proteins and cognition (memory and executive function). As in the ROSMAP analysis, we used a logistic regression with subtype (coded as before) as the dependent variable and demographic, cognitive and CSF-based measures as the explanatory variables. As only a subset of subjects had measures of sTREM2 and progranulin, we performed two separate regression analyses, testing for: (i) differences in demographics, stage, cognition and CSF total tau and amyloid beta in 259 subjects and (ii) demographics, stage, sTREM2 and progranulin in 190 subjects.

## Data availability

The post mortem immunohistochemistry data used in this study comes from the Religious Orders Study (ROS) and Rush Memory and Aging Project (MAP) studies, both of which can be obtained from the Rush Alzheimer’s Disease Center (RADC) by submitting a data request via the https://www.radc.rush.edu website. The in vivo PET and CSF data used in this study comes from the Alzheimer’s Disease Neuroimaging Initiative (ADNI) database (adni.loni.usc.edu).

## Code availability

A python-based implementation of SuStaIn (pySuStaIn), supporting both z-score style and mixture style data likelihoods, is available at https://github.com/ucl-pond/pySuStaIn. The kernel density estimation-based mixture modeling code is available at https://github.com/noxtoby/kde_ebm_open.

## Acknowledgements

This project has received funding from the European Union’s Horizon 2020 research and innovation program under grant agreement No. 666992. NPO is a UKRI Future Leaders Fellow (MRC MR/S03546X/1). This project was supported by the National Institute for Health Research University College London Hospitals Biomedical Research Centre. PAW was funded by a Medical Research Council Skills Development Fellowship (MR/T027770/1). ALY is supported by an MRC Skills Development Fellowship (MR/T027800/1).

Data collection and sharing for this project was funded by the Alzheimer’s Disease Neuroimaging Initiative (ADNI) (National Institutes of Health Grant U01 AG024904) and DOD ADNI (Department of Defense award number W81XWH-12-2-0012). ADNI is funded by the National Institute on Aging, the National Institute of Biomedical Imaging and Bioengineering, and through generous contributions from the following: AbbVie, Alzheimer’s Association; Alzheimer’s Drug Discovery Foundation; Araclon Biotech; BioClinica, Inc.; Biogen; Bristol-Myers Squibb Company; CereSpir, Inc.; Cogstate; Eisai Inc.; Elan Pharmaceuticals, Inc.; Eli Lilly and Company; EuroImmun; F. Hoffmann-La Roche Ltd and its affiliated company Genentech, Inc.; Fujirebio; GE Healthcare; IXICO Ltd.; Janssen Alzheimer Immunotherapy Research & Development, LLC.; Johnson & Johnson Pharmaceutical Research & Development LLC.; Lumosity; Lundbeck; Merck & Co., Inc.; Meso Scale Diagnostics, LLC.; NeuroRx Research; Neurotrack Technologies; Novartis Pharmaceuticals Corporation; Pfizer Inc.; Piramal Imaging; Servier; Takeda Pharmaceutical Company; and Transition Therapeutics. The Canadian Institutes of Health Research is providing funds to support ADNI clinical sites in Canada. Private sector contributions are facilitated by the Foundation for the National Institutes of Health (www.fnih.org). The grantee organization is the Northern California Institute for Research and Education, and the study is coordinated by the Alzheimer’s Therapeutic Research Institute at the University of Southern California. ADNI data are disseminated by the Laboratory for Neuro Imaging at the University of Southern California.

