## Supplementary Material for "Tau-first subtype of Alzheimer’s disease consistently identified across in vivo and post mortem studies"

**Supplementary Figure 1**

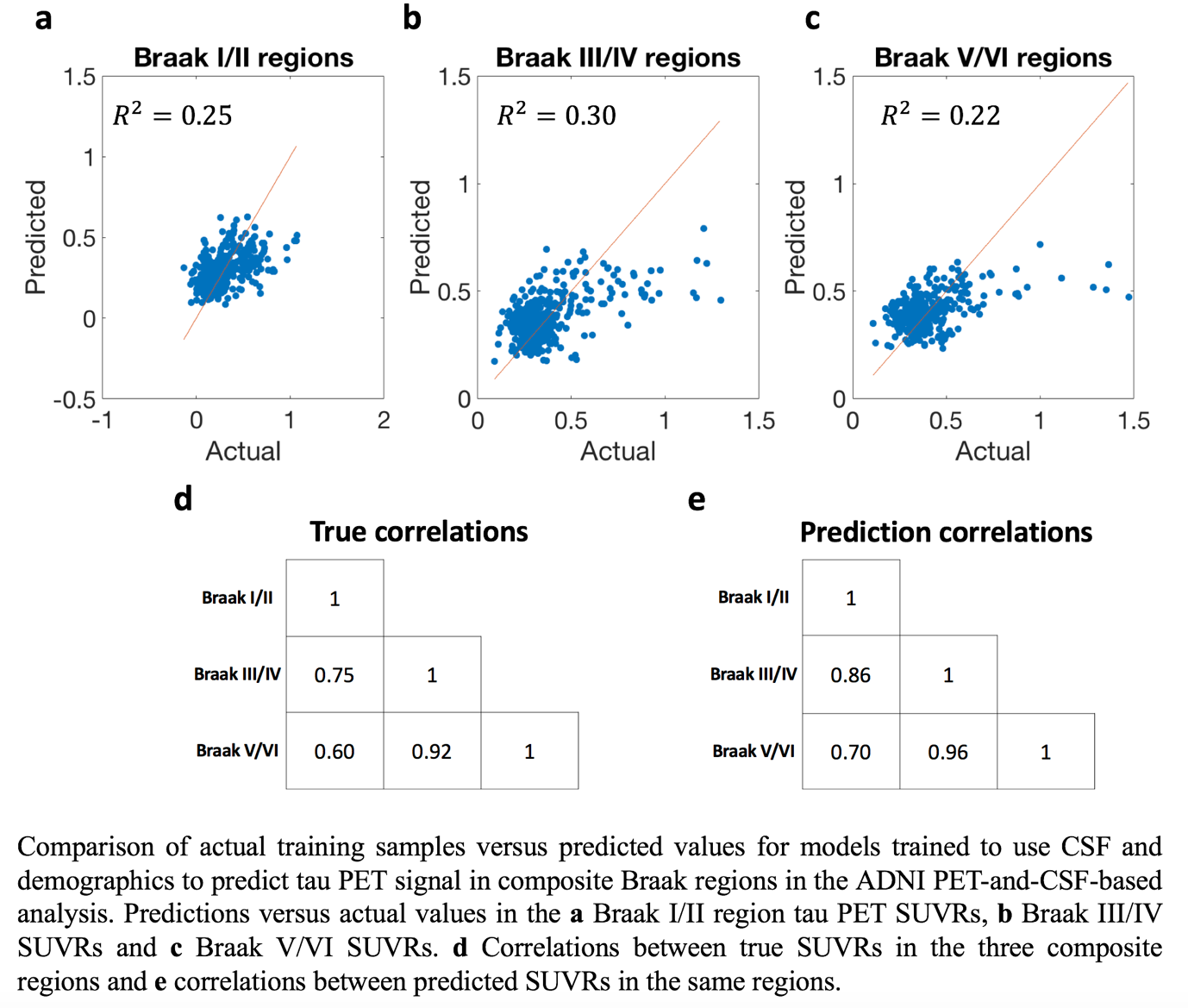

**Supplementary Figure 2**

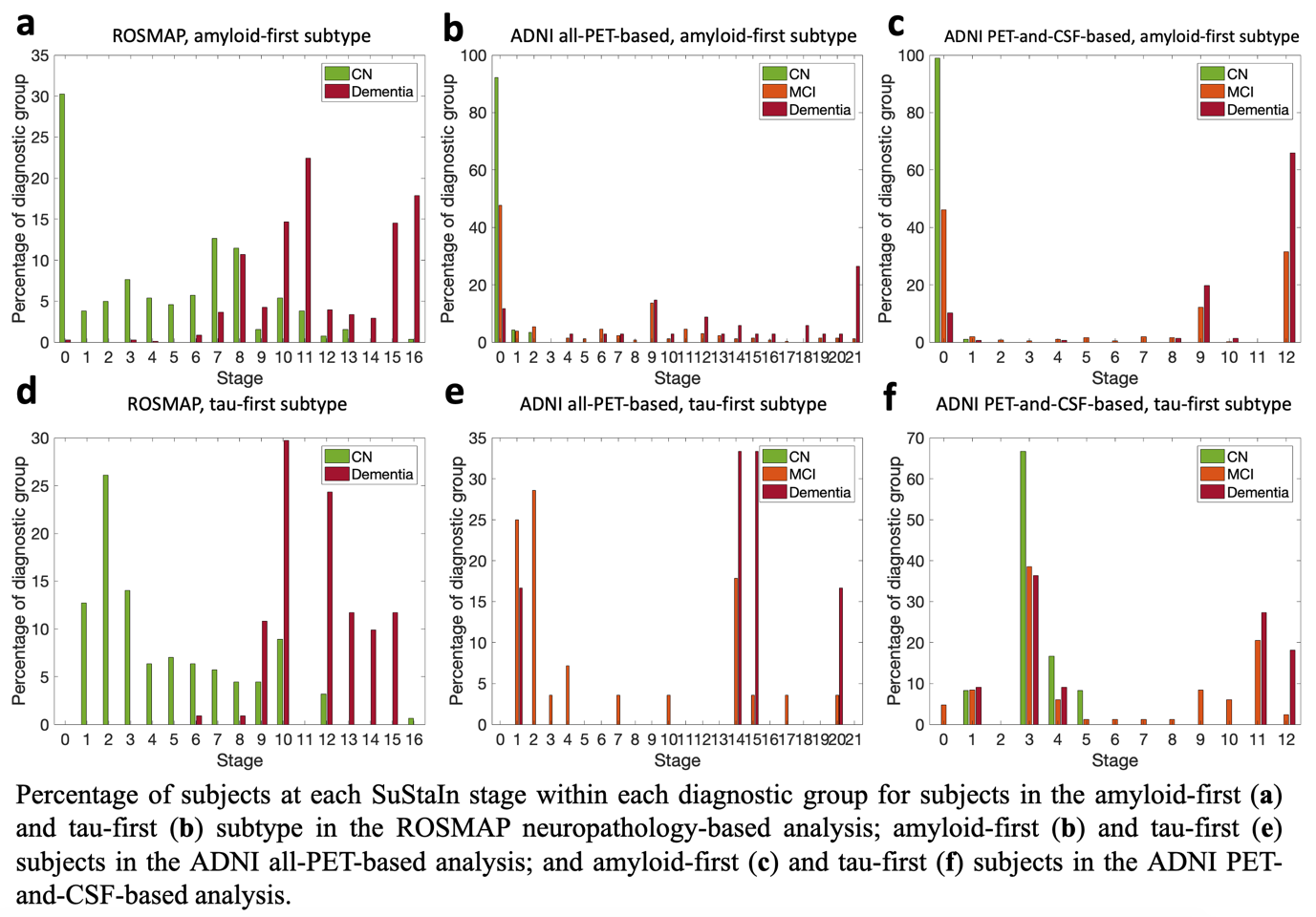

**Supplementary Table 1**

| Amyloid PET regions | |
| --- | --- |
| Composite SUVR | Component SUVRs |
| Cingulate | Caudal Anterior Cingulate  Rostral Anterior Cingulate  Posterior Cingulate  Isthmus Cingulate |
| Inferior Frontal | Pars Opercularis  Pars Orbitalis  Pars Triangularis |
| Middle Frontal | Caudal Middle Frontal  Rostral Middle Frontal |
| Superior Frontal | Superior Frontal |
| Orbitofrontal | Lateral Orbitofrontal  Medial Orbitofrontal |
| Precuneus | Precuneus |
| Lateral Parietal | Inferior Parietal  Superior Parietal  Supramarginal  Post Central |
| Lateral Temporal | Inferior Temporal  Middle Temporal  Superior Temporal  Transverse Temporal |
| Medial Temporal | Entorhinal  Parahippocampal  Fusiform |
| Tau PET regions | |
| Composite SUVR | Component SUVRs |
| Amygdala | Amygdala |
| Entorhinal | Entorhinal |
| Cingulate | Caudal Anterior Cingulate  Rostral Anterior Cingulate  Posterior Cingulate  Isthmus Cingulate |
| Medial Temporal | Parahippocampal  Fusiform |
| Lateral Temporal | Inferior Temporal  Middle Temporal  Superior Temporal  Transverse Temporal |
| Inferior Frontal | Pars Opercularis  Pars Orbitalis  Pars Triangularis |
| Middle Frontal | Caudal Middle Frontal  Rostral Middle Frontal |
| Superior Frontal | Superior Frontal |
| Orbitofrontal | Lateral Orbitofrontal  Medial Orbitofrontal |
| Lateral Parietal | Inferior Parietal  Superior Parietal  Supramarginal  Post Central |
| Precuneus | Precuneus |
| Occipital | Cuneus  Lateral Occipital  Lingual  Pericalcarine |

Composite SUVRs used in our analysis and their component, Freesurfer-based regional SUVRs.

**Supplementary Table 2**

|  | **ROSMAP** | **ADNI tau-PET** | **ADNI tau-CSF** | **ADNI Comparison** |
| --- | --- | --- | --- | --- |
| n | 843 | 197 | 261 | N/A |
| Age, Mean $\pm$ SD | 89.9 $\pm$ 6.4 | 77.1 $\pm$ 7.6 | 73.8 $\pm$ 7.2 | 3.42 × 10^-6^ * |
| Years since baseline, Mean $\pm$ SD | N/A | 4.0 $\pm$ 3.6 | 0.4 $\pm$ 1.3 | < 1 × 10^-6^ * |
| Education Years, Mean $\pm$SD | 15.9 $\pm$ 3.5 | 16.4 $\pm$ 2.6 | 16.4 $\pm$ 2.7 | 0.83 |
| Females, Percentage | 71% | 49% | 53% | 0.34 |
| APOE4 alleles (% 0,1,2) | N/A | 51%, 37%, 12% | 47%, 43%, 10% | 0.42 |
| Diagnosis (% CN,MCI,AD) | 39%, 0%, 61% | 5%, 82%, 14% | 5%, 78%, 17% | 0.58 |

Characterization of cohorts used in subtype analyses. We excluded one out of 198 subjects from the ADNI tau-PET based cohort due to missing age. We compared the ADNI tau-PET and ADNI tau-CSF cohorts via statistical tests: (i) age, years since study baseline and education years via one-way ANOVAs; (ii) percentage female, percentage with 0,1 or 2 APOE4 alleles and percentage in each diagnostic category (CN/MCI/AD) via chi-squared tests. P-values of these tests are reported in right-hand column. * P < 0.05

**Supplementary Table 3**

|  | Estimate | SE | t-statistic | P-value | Odds Ratio |
| --- | --- | --- | --- | --- | --- |
| Intercept | -1.24 | 0.37 | N/A | N/A | N/A |
| Age | 0.11 | 0.08 | 1.33 | 0.18 | 1.12 |
| Sex | -0.04 | 0.08 | -0.50 | 0.62 | 0.96 |
| Education Years | 0.03 | 0.02 | 1.14 | 0.25 | 1.03 |
| Stage | -0.51 | 0.09 | -5.76 | < 1 × 10^-6^ * | 0.60 |
| TDP-43 Stage | 0.16 | 0.09 | 1.75 | 0.08 | 1.17 |
| CAA Stage | -0.09 | 0.08 | -1.09 | 0.28 | 0.91 |
| Cerebral Atherosclerosis Stage | -0.06 | 0.09 | -0.67 | 0.51 | 0.94 |
| Arteriolosclerosis Stage | 0.13 | 0.08 | 1.48 | 0.14 | 1.13 |
| Hippocampal Sclerosis | -0.08 | 0.09 | -0.89 | 0.37 | 0.92 |

Logistic regression-based analysis of 810 subjects’ (554 amyloid-first, 256 tau-first) demographic and comorbid pathology differences between subtypes in the ROSMAP neuropathology-based analysis. * P < 0.05

**Supplementary Table 4**

|  | Estimate | SE | t-statistic | P-value | Odds Ratio |
| --- | --- | --- | --- | --- | --- |
| Intercept | -0.69 | 0.18 | -3.74 | 1.86 × 10^-4^ * | N/A |
| Age | -0.56 | 0.24 | -2.32 | 0.02 * | 0.57 |
| Sex | -0.36 | 0.21 | -1.69 | 0.09 | 0.70 |
| APOE4 | -0.16 | 0.22 | -0.76 | 0.45 | 0.85 |
| Education Years | 0.10 | 0.19 | 0.54 | 0.59 | 1.11 |
| Stage | 0.21 | 0.24 | 0.88 | 0.38 | 1.24 |
| CSF Tau | 1.97 | 0.31 | 6.33 | < 1 × 10^-6^ * | 7.18 |
| CSF Amyloid Beta | 1.04 | 0.31 | 3.37 | 7.53 × 10^-4^ * | 2.83 |
| Memory | 0.30 | 0.3 | 0.99 | 0.32 | 1.35 |
| Executive Function | -0.10 | 0.29 | -0.35 | 0.73 | 0.90 |

Logistic regression-based analysis of subtype differences in 259 subjects’ (161 amyloid-first, 98 tau-first) demographics, stage, CSF-based amyloid and tau measures, memory (ADNI-MEM) and executive function (ADNI-EF) in the ADNI tau-CSF-based analysis. * P < 0.05

**Supplementary Table 5**

|  | Estimate | SE | t-statistic | P-value | Odds Ratio |
| --- | --- | --- | --- | --- | --- |
| Intercept | -0.56 | 0.17 | -3.19 | 1.44 × 10^-3^ * | N/A |
| Age | -0.57 | 0.19 | -2.99 | 2.83 × 10^-3^ * | 0.56 |
| Sex | 0.13 | 0.19 | 0.70 | 0.49 | 1.14 |
| APOE4 | 0.12 | 0.19 | 0.61 | 0.54 | 1.12 |
| Education Years | 0.11 | 0.19 | 0.59 | 0.56 | 1.12 |
| Stage | -0.38 | 0.19 | -2.03 | 0.04 * | 0.68 |
| sTREM2 | 0.89 | 0.21 | 4.18 | 2.92 × 10^-5^ * | 2.43 |
| Progranulin | 0.28 | 0.21 | 1.34 | 0.18 | 1.32 |

Logistic regression-based analysis of subtype differences in 190 subjects’ (116 amyloid-first, 74 tau-first) demographics, stage and CSF-based sTREM2 and progranulin differences in the ADNI tau-CSF-based analysis. * P < 0.05
